# Rieske head domain dynamics and indazole-derivative inhibition of *Candida albicans* complex III

**DOI:** 10.1101/2021.05.18.444650

**Authors:** Justin M. Di Trani, Zhongle Liu, Luke Whitesell, Peter Brzezinski, Leah E. Cowen, John L. Rubinstein

## Abstract

During cellular respiration, electron transfer between the integral membrane protein complexes of the electron transport chain is coupled to proton translocation across the inner mitochondrial membrane, which in turn powers synthesis of ATP and transmembrane transport processes. The homodimeric electron transport chain Complex III (CIII_2_) oxidizes ubiquinol (UQH_2_) to ubiquinone (UQ), transferring electrons to cytochrome *c*, and translocating protons through a mechanism known as the Q cycle. The Q cycle involves UQH_2_ oxidation and UQ reduction at two different sites within each CIII monomer, as well as movement of the head domain of the Rieske subunit. We used cryoEM to determine the structure of CIII_2_ from *Candida albicans*, revealing density for endogenous UQ in the structure and allowing us to directly visualize the continuum of conformations of the Rieske head domain. Analysis of these conformations does not indicate cooperativity in the position of the Rieske head domains or binding of ligands in the two CIIIs of the CIII_2_ dimer. CryoEM with the indazole derivative Inz-5, which inhibits fungal CIII_2_ and is fungicidal when administered with fungistatic azole drugs, showed that inhibition by Inz-5 alters the equilibrium of the Rieske head domain positions.

## INTRODUCTION

Aerobic organisms use energy released by the oxidation of nutrients to synthesize the chemical energy currency adenosine triphosphate (ATP) from adenosine diphosphate (ADP) and inorganic phosphate. This process, known as oxidative phosphorylation, may be targeted by small molecules that serve as antibiotics (Andries *et al*., 2005) and antifungals (Roemer and Krysan, 2014). The yeast *Candida albicans* is an opportunistic pathogen that most often infects immunocompromised patients, which can lead to organ failure and death (Pfaller and Diekema, 2010). The limited arsenal of effective and well-tolerated antifungal drugs (Roemer and Krysan, 2014) and the emergence of resistance to these therapeutics (Mazu *et al*., 2016) has highlighted the need for development of new antifungals. High-throughput screening led to identification of a series of indazole derivatives that target oxidative phosphorylation and can be administered with fungistatic azole compounds converting them into fungicidal drugs. One of these derivatives, Inz-5, showed sub-micromolar activity against *C. albicans*, favourable selectivity over human cells, and was less susceptible to metabolic inactivation than other derivatives (Vincent *et al*., 2016).

During oxidative phosphorylation, oxidation of carbon sources and ATP synthesis are coupled energetically by the formation of an electrochemical transmembrane proton motive force (PMF). A group of membrane protein complexes, known collectively as the respiratory chain or electron transport chain, generate the PMF by translocating protons from the negative (N) side to the positive (P) side of the membrane. Mitochondrial respiratory chains make use of a hydrophobic ubiquinone (UQ) in the membrane and soluble cytochrome *c* (cyt. *c*) in the intermembrane space as intermediate electron carriers between the membrane protein complexes. In *C. albicans* UQ can be reduced by either NADH:UQ oxidoreductase (Complex I, CI) or succinate:UQ oxidoreductase (Complex II, CII). The resultant ubiquinol (UQH_2_) is the electron donor for UQH_2_:cyt. *c* oxidoreductase (Complex III, CIII), also known as cyt. *bc*_1_, an obligate homodimeric assembly that serves as the penultimate electron-transport complex. The antifungal indazole derivatives described above target the *C. albicans* CIII in a fungal-selective manner (Vincent *et al*., 2016).

CIII contributes to the PMF through a mechanism known as the Q cycle (for reviews see (Xia *et al*., 2013; Sarewicz and Osyczka, 2015)). Briefly, the cyt. *b* subunit of CIII possesses two UQ binding sites: one near the P side of the membrane (the Q_P_ site) and the other near the N side of the membrane (the Q_N_ site). During catalysis, UQH_2_ binds at the Q_P_ site where it is oxidized, first to semiquinone (UQ^•-^) and then to UQ. This oxidation releases two protons from UQH_2_ to the P side of the membrane. The first electron from UQH_2_ reduces an iron sulphur (FeS) cluster in a mobile head domain of the Rieske protein subunit of CIII (Zhang *et al*., 1998; Tian *et al*., 1999; Xia, Yu and Yu, 2000; Darrouzet *et al*., 2001). The second electron from UQH_2_ is transferred consecutively to heme *b*_L_, heme *b*_H_, and a UQ molecule bound at the Q_N_ site, reducing it to UQ^•-^ (Wikström and Berden, 1972). The Rieske protein head domain moves from its position near the Q_P_ site (known as its *b* position due to its proximity to heme *b*_L_) to a position near the heme of cyt. *c*_1_ (known as its *c* position). In the *c* position, the electron is transferred from FeS to cyt. *c*_1_ and then to water-soluble cyt. *c*, which diffuses to cytochrome *c* oxidase (Complex IV, CIV), also known as cyt. *aa*_3_, and catalyzes reduction of molecular oxygen to water. The path taken by the first electron from UQH_2_ in the Q_P_ site is known as the *c* chain due to the involvement of heme *c*_1_ and cyt. *c*, while the path of the second electron is known as the *b* chain due to the involvement of hemes *b*_L_ and *b*_H_. Repetition of this sequence of events from a second UQH_2_ binding at the Q_P_ site leads to release of two more protons to the P side of the membrane, transfer of a second electron to CIV, and reduction of UQ^•-^ to UQH_2_ at the Q_N_ site, which is associated with uptake of two protons from the N side of the membrane.

A number of Q_P_ site inhibitors are known to bind competitively with UQH_2_ and have been used for medical and agricultural applications (Esser *et al*., 2004). Crystal structures of CIII_2_ have been determined in complex with several of these inhibitors (Kim *et al*., 1998; Tian *et al*., 1999; Esser *et al*., 2006; Berry and Huang, 2011). Inhibitor binding at the Q_P_ site alters the position of the Rieske head domain in different ways depending on the inhibitor pose in the Q_P_ pocket and whether the inhibitor forms direct contacts with the Rieske head domain (Kim *et al*., 1998; Esser *et al*., 2004). This influence on position has allowed for classification of Q_P_ site inhibitors into two classes. With the first class, the Rieske head domain remains mobile (P_m_ type inhibitors), while the second class of inhibitors fix the Rieske head in the *b* position (P_f_ type inhibitors). P_m_ type inhibitors includes azoxystrobin and other compounds that contain the β-methoxyacrylate (MOA) moiety (Esser *et al*., 2004, 2019). Inhibitors containing the MOA moiety are referred to as strobilurins, a family of widely utilized agricultural fungicides (Bartlett *et al*., 2002; Zhang *et al*., 2020). The antibiotic myxothiazol is also a P_m_ type inhibitor (Thierbach and Reichenbach, 1981). In contrast, P_f_ type inhibitors include UQ analogues such as 5-undecyl-6-hydroxy-4,7-dioxobenzothiazole (UHDBT) and stigmatellin (Iwata *et al*., 1998; Esser *et al*., 2004), the latter of which forms a hydrogen bond with the Rieske head domain to hold it in the *b* position (Zhang *et al*., 1998). A similar interaction between the Rieske protein and substrate UQH_2_ has been hypothesized to stabilize the Rieske protein head domain in the *b* position (Crofts and Wang, 1989; Link and Iwata, 1996; Jünemann, Heathcote and Rich, 1998; Millett *et al*., 2013; Scharlau *et al*., 2019). This interaction underpins models where the Rieske head domain movement is correlated with UQH_2_ binding in order to ensure bifurcation of electron transfer along the *c* and *b* chains of the Q cycle (Berry and Huang, 2003; Sarewicz and Osyczka, 2015). Alternative models suggest that the Rieske head domain moves stochastically and concerted motion is not required for maintaining the Q cycle (for review see (Sarewicz and Osyczka, 2015)). Furthermore, data suggest a link between ligand binding at the Q_P_ sites and the dynamics of the Rieske head domains of individual monomers (Covian, Gutierrez-Cirlos and Trumpower, 2004; Covian *et al*., 2007; Covian and Trumpower, 2008) and were also interpreted as showing anti-cooperative substrate binding in the Q_P_ sites in the two halves of the CIII_2_ dimer (Covian *et al*., 2007).

Here we determined the structure of CIII_2_ from *C. albicans* by electron cryomicroscopy (cryoEM). While crystallization for X-ray diffraction tends to select a single protein conformation, cryoEM enables analysis of multiple conformations that may exist simultaneously in solution (e.g. (Zhao, Benlekbir and Rubinstein, 2015; Huang *et al*., 2016)). Probabilistic principal component analysis (PPCA) (Punjani and Fleet, 2020) revealed three positions for the Rieske head domain: the *b* position, *c* position, and an intermediate position. Comparison of the monomers in the CIII_2_ dimer indicates that the Rieske subunits act independently, arguing against cooperativity in the two halves of the CIII_2_ dimer. Density for UQ in the Q_P_ site in the cryoEM maps, in a position similar to stigmatellin (Zhang *et al*., 1998), indicates that the hydrogen bond seen between stigmatellin and the Rieske head domain can occur with the enzyme’s natural substrate, providing support for models of the Q cycle that involve hydrogen bond mediated electron and proton transfer between UQH_2_ and the Rieske head domain. We also determined the structure of CIII_2_ with Inz-5 bound within the Q_P_ site, which explains the compound’s specificity for fungal CIII. The binding pose of Inz-5 is similar to that of strobilurin family of inhibitors. PPCA of the Rieske head domain shows that Inz-5 is a P_m_ type inhibitor that mobilizes the Rieske head domain, allowing it to shift from the *b* to the *c* position. These similarities to strobilurins and previous studies in fungi suggest that Inz-5 and other indazole derivatives may serve as lead compounds that could be optimized for use as agricultural fungicides or to treat fungal infections in humans.

## RESULTS

### Candida CIII_2_ overall structure

DNA sequence encoding a His_6_-3×FLAG tag was introduced into the *C. albicans* genome immediately downstream of both alleles of the gene encoding the Qcr2 subunit of CIII. Following growth in a rich medium with galactose as the carbon source, cells were collected, their mitochondria harvested, mitochondrial membranes isolated and solubilized with detergent, and CIII_2_ purified by affinity chromatography. SDS-PAGE showed bands for eight of the complex’s nine subunits (**Fig. 1A**), with the small hydrophobic subunit Qcr9 being difficult to detect. UV-visible spectroscopy showed the presence of the expected hemes in the preparation (**Fig. S1A)**.

**Fig. 1.**
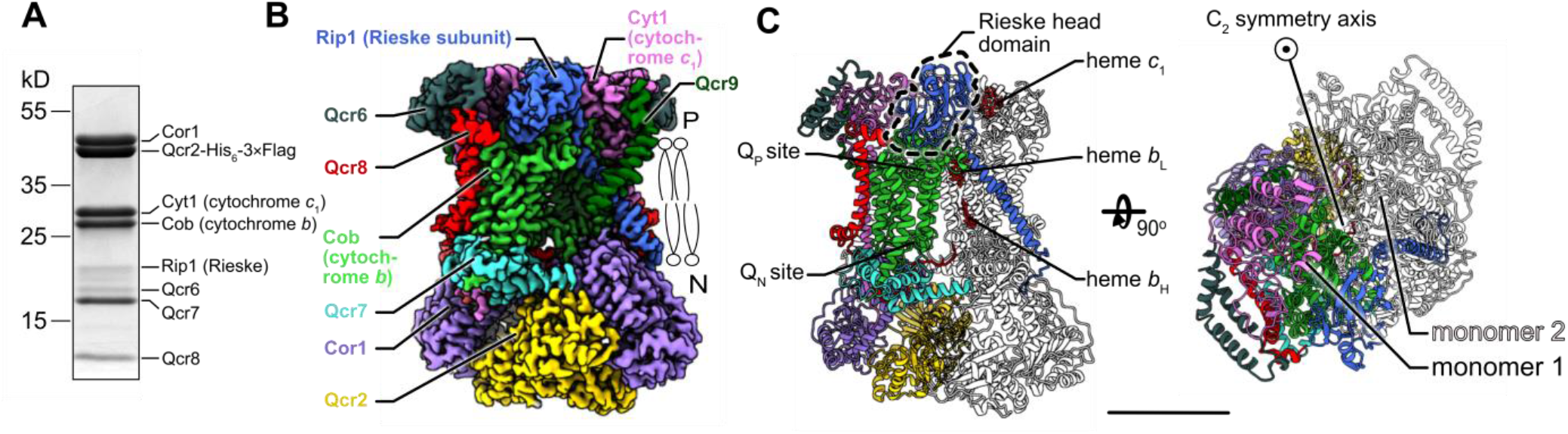
Overall structure of *C. albicans* CIII_2_. (**A**) SDS-PAGE, (**B**) cryoEM map, and (**C**) atomic model of *C. albicans* CIII_2_. Scale bar, 50 Å.

CryoEM of the preparation led to a map with an overall resolution of 3.0 Å (**Fig. 1B and S1B-E**). Map resolution for the Rieske head domain was ∼4 to 6 Å (**Fig. S1E**), preventing *de novo* construction of an atomic model for this part of the complex but allowing for rigid body fitting of a homology model based on the structure of the *Saccharomyces cerevisiae* CIII Rieske head domain (Hunte *et al*., 2000). The lower resolution in this region of the map suggests variability in the position of the Rieske head domain. Map resolution for the rest of the complex enabled construction of an atomic model for almost all residues except for residues 211 to 220 in subunit Qcr2, 40 to 44 in Qcr8, and seven other residues throughout the complex, all of which were modelled as alanine **(Fig. 1C, S1F and Table S1**,**2**).

Density for endogenous UQ in the CIII_2_ dimer is found in the two Q_P_ sites (**Fig. S2A**) and two Q_N_ sites (**Fig. S2B**). Importantly, the density for UQ in the Q_P_ and Q_N_ sites was lower than for adjacent parts of the protein, suggesting a low occupancy of UQ in these sites. The position occupied by UQ at the Q_P_ site is similar to the previously-observed position for the CIII inhibitor stigmatellin, the aforementioned UQ analogue that forms a hydrogen bond with the Rieske head domain (**Fig. S2C**) (Zhang *et al*., 1998). The Q_P_ binding site is large and may accommodate UQ in a number of different orientations (Brandt and von Jagow, 1991; Crofts, Hong, *et al*., 1999; Hong *et al*., 1999). However, the orientation found for UQ in the Q_P_ site in the current study, as well as in a recent cryoEM map of the ovine CICIII_2_ supercomplex, supports models in which the Rieske head domain forms a hydrogen bond with substrate to receive the first electron and proton from UQH_2_ (Berry and Huang, 2003; Sarewicz and Osyczka, 2015).

### Rieske protein appears in three positions

To investigate the variability of the Rieske head domain position in the CIII_2_ structure, we performed three-dimensional variability analysis (3DVA) (Punjani and Fleet, 2020), which makes use of probabilistic principal component analysis (PPCA) to separate particle images that contribute to 3D maps of flexible protein structures. The analysis assigns each particle image a coefficient along components that describe motions as linear additions of density to a consensus structure (**Fig. S3**). Following symmetry expansion to allow inclusion of both CIII monomers from each CIII_2_ dimer, 3DVA (**Fig. S3A**) revealed three Rieske head domain positions, with 3D refinement used to improve density for the Rieske head domain in maps from the different clusters of particle images. In the first position, the FeS cluster of the Rieske head domain is close to the Q_P_ site (**Fig. 2 left**), in the second the FeS cluster is close to the *c*_1_ heme (**Fig. 2 right**), while the third position shows the Rieske head domain between these two states (**Fig. 2 middle** and **Movie 1**). While distinct Rieske head domain positions have been observed previously in independent crystal structures, the analysis presented here shows that a single population of CIII_2_ complexes in solution possesses a distribution of head domain positions.

**Fig. 2.**
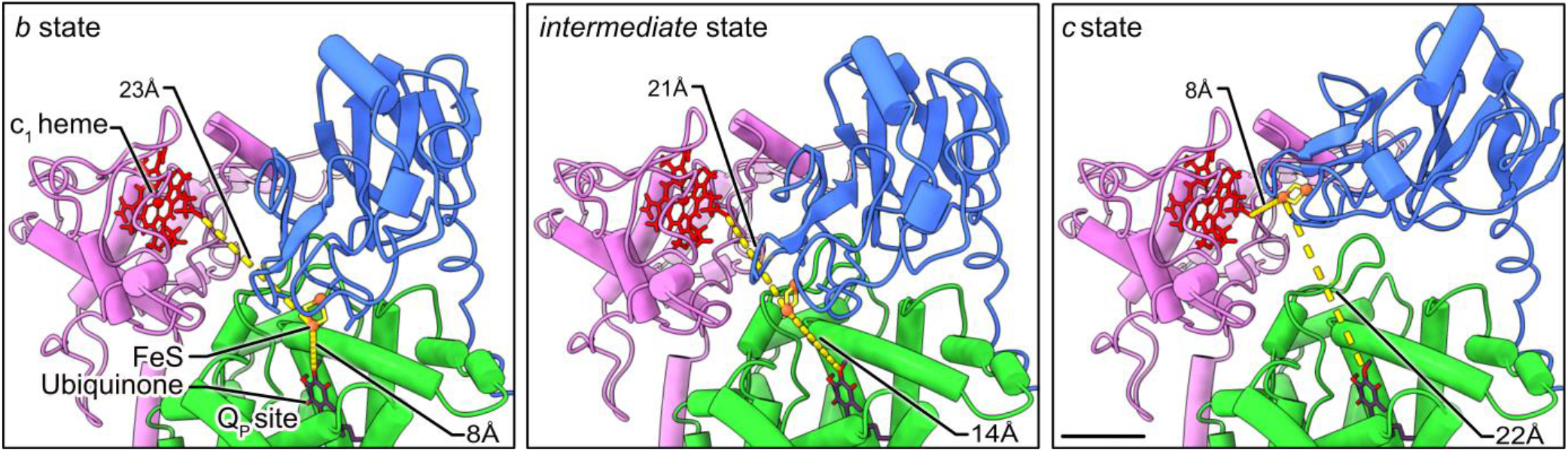
The Rieske head domain is found in three positions. Rigid body fit of Rieske head domain backbone and FeS cluster into maps refined from the *b, intermediate*, and *c* state particle images. Scale bar, 10 Å.

With the Rieske head domain near the Q_P_ site, the FeS cluster is 8 Å from UQ (edge to edge), similar to the distance between the FeS cluster and stigmatellin in a stigmatellin-bound crystal structure of CIII_2_ (Crofts, Guergova-Kuras, *et al*., 1999; Osyczka, Moser and Dutton, 2005). This orientation of the Rieske head domain matches crystal structures with the domain in the *b* position, with Cα RMSDs of 3.2 and 3.6 Å from bovine and *S. cerevisiae* structures, respectively (Lange and Hunte, 2002; Huang *et al*., 2005). With the Rieske head domain between the Q_P_ site and cyt. *c*_1_, the FeS cluster is 14 Å from the Q_P_ site and 21 Å from the edge of the heme. This conformation matches the intermediate state observed previously in a crystal structure of CIII in the absence of inhibitors, with a Cα RMSD of 2.6 Å for Rieske head domains (Iwata *et al*., 1998). With the Rieske head domain close to cyt. *c*_1_, the edge of the FeS cluster is 8 Å from the edge of the heme, matching the *c* position from a crystal structure with the Q_P_ site inhibitor myxothiazol and Q_N_ site inhibitor antimycin (Cα RMSD of 2.9 Å for Rieske head domains) (Iwata *et al*., 1998). Altogether, the FeS cluster spans a distance of ∼17 Å between the *b* and *c* positions. It is important to note that the observation of three states here does not preclude the existence of additional intermediate states seen previously in crystallographic studies (Esser *et al*., 2019).

### Inz-5 binds to the Q_P_ site of CIII_2_

Small molecules that bind at the Q_P_ site can alter the Rieske head position (Kim *et al*., 1998; Berry and Huang, 2011) and we investigated the possibility of this effect with Inz-5 (**Fig. 3A**). Previous mutational analysis suggests that Inz-5 inhibits CIII by binding in the Q_P_ site (Pfaller and Diekema, 2010; Vincent *et al*., 2016). In UQH_2_:cyt. *c* oxidoreductase assays Inz-5 inhibits CIII_2_ with an IC_50_ of 24 ± 3 nM (mean ± s.d., *n*=4 measurements per inhibitor concentration) (**Fig. 3B**), ∼10× lower than the IC_50_ measured with mitochondria using a slightly lower concentration of the same substrate (Vincent *et al*., 2016). In order to elucidate how Inz-5 binds the enzyme, the inhibitor was added to purified CIII_2_ at a 1.8-fold molar excess relative to CIII monomers, cryoEM specimens were prepared, and the protein-inhibitor complex was subjected to structure determination. The resulting map of CIII_2_ with Inz-5 bound was refined to 3.3 Å resolution (**Fig. S1C**) and shows strong density for Inz-5 in the Q_P_ sites (**Fig. 3C**) with no detectable Inz-5 density in the Q_N_ sites. The Rieske head domain was subject to the same classification scheme used for images of CIII_2_ without Inz-5.

**Fig. 3.**
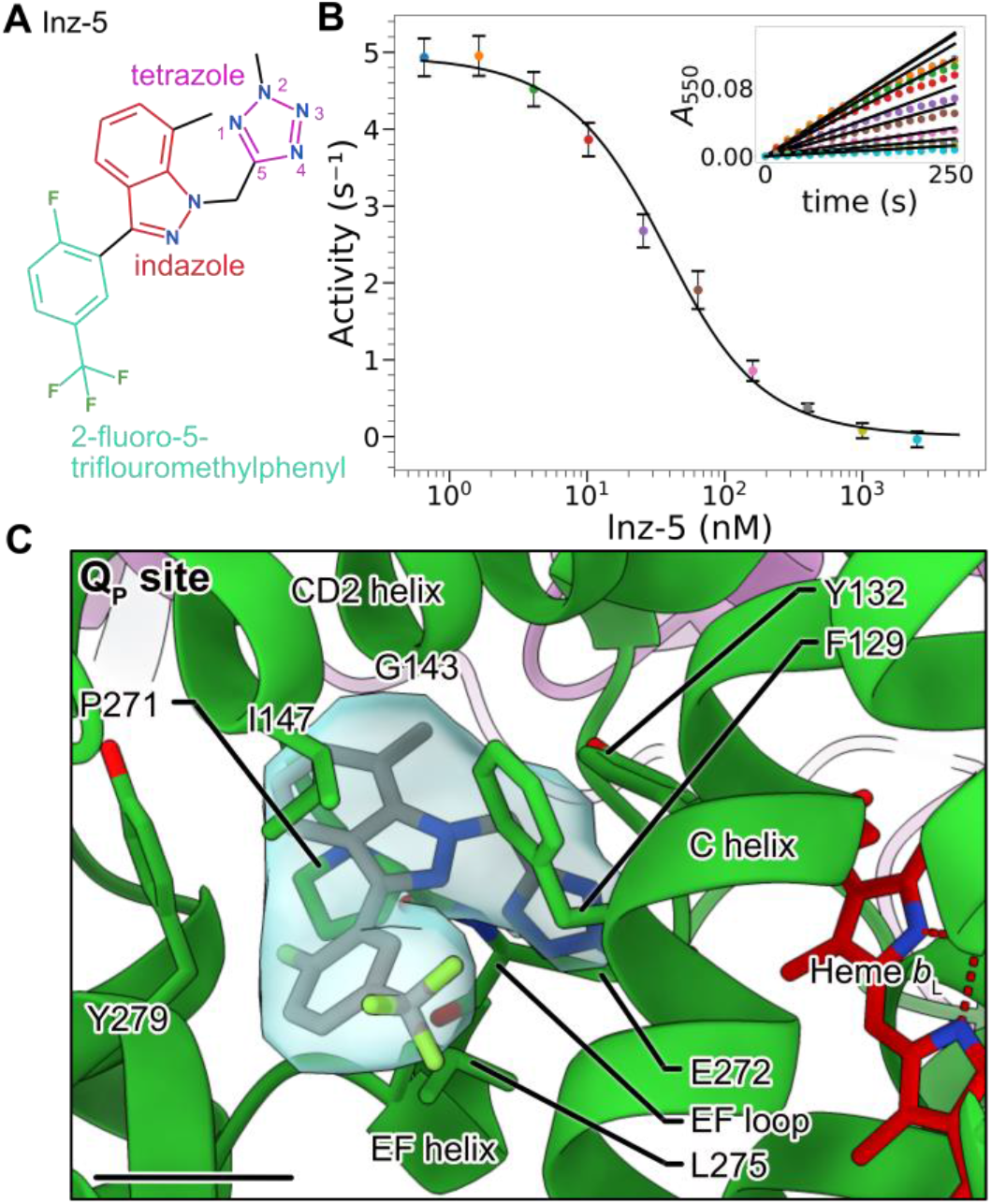
Inhibition of *C. albicans* CIII_2_ by Inz-5. (**A**) Structure of Inz-5. (**B**) Inhibition curve for decylubiquinol (DBH_2_):cyt. *c* oxidoreductase activity of CIII_2_ upon incubation with Inz-5. Data is presented as mean ± s.d. (*n*=4 independent measurements), solid black line is fit of data. The inset shows representative raw data (points) and initial rate linear fits (black lines). (**C**) Binding environment of Inz-5 in the Q_P_ site with α helices, loops, and key residues labelled. Scale bar, 5 Å.

Inz-5 in the Q_P_ binding pocket is enclosed by the transmembrane C helix, the EF helix, and the EF loop, which are adjacent to heme *b*_L_, and the CD2 helix (**Fig. 3C** and **Movie 2**). Importantly, the EF helix, EF loop, and CD2 helix also interact with the Rieske head domain when in the *b* state (Esser *et al*., 2006; Berry and Huang, 2011). Inz-5 wraps around the EF helix with the tetrazole moiety pointing toward the *b*_L_ heme and the 2-fluoro-5-triflouromethylphenyl moiety pointing toward the membrane accessible side of the Q_P_ site. Compared to the inhibitor-free structure, Glu272 of cyt. *b* changes orientation in order to accommodate the tetrazole moiety (**Fig. S4A**). Overall, the binding pose of Inz-5 is similar to that of the strobilurin fungicide azoxystrobin, which has been crystallized with CIII_2_ from *Rhodobacter sphaeroides* (Esser *et al*., 2019), and β-methoxyacrylate stilbene (MOAS) crystallized with bovine CIII_2_ (Esser *et al*., 2004). Many of the stabilizing interactions that occur in Inz-5 are also apparent in the strobilurin-bound structures. The tetrazole, indazole, and 2-fluoro-5-triflouromethylphenyl trifluoride moieties of Inz-5 are each ∼5 Å from Tyr132, Tyr279, and Phe129 of the cyt. *b* subunit, respectively, allowing for the formation of stabilizing aromatic pairs (Burley and Petsko, 1985). Similarly, the phenyl group of azoxystrobin and α-benzene of MOAS both form aromatic pairs with Tyr279 of cyt. *b* (equivalent to Tyr278 in *R. sphaeroides* and bovine CIII). In addition, the pyrimidine moiety of azoxystrobin and terminal benzene of the stilbene moiety form aromatic pairs with Phe129 of cyt. *b* (Phe128 in *R. sphaeroides* and *bovine* CIII) (**Fig. S4B**,**C**). In the current structure, it appears that N4 of the tetrazole ring is sufficiently close to the backbone amide of Glu272 from cyt. *b* (Glu271 in *R. sphaeroides* and bovine CIII) for formation of a hydrogen bond (**Fig. 3C**). This amide also forms a hydrogen bond with the methyl ester oxygen of the MOA group of azoxystrobin (**Fig. S4B-C**). Similar to the α-benzene of MOAS and the phenyl group of azoxystrobin, the indazole group of Inz-5 is wedged between Pro271, Gly143, and Leu275 of cyt. *b* (Pro270, Gly142, and Phe274, respectively in bovine and *R. sphaeroides* CIII).

### Inz-5 is a P_m_ type inhibitor

As described above, the binding pose of an inhibitor can influence the Rieske head domain position. To assess the effect of Inz-5 on Rieske head domain position, particle images with and without the compound were combined and subjected to 3DVA with a mask that included only the Rieske head domain. Combining the datasets prior to 3DVA is necessary to ensure that the algorithm identifies the same principal components for the two datasets. This analysis separates the particle images into populations showing the Rieske head domain in the *b, intermediate*, and *c* positions. Further 3DVA with the combined *b* and *intermediate* populations, as well as the combined *c* and *intermediate* populations, yielded components that show structural changes between the *b* and *intermediate* states, and between the *intermediate* and *c* states (**Fig. 4A,B**), and assigned coefficients to particle images along these components. A distinct separation of populations along these principal components is apparent for the compound-free and Inz-5-bound datasets (**Fig. 4C,D**, light and dark blue, respectively), with the Inz-5-bound particle images assigned mostly to the *c* state and the compound-free particle images assigned more evenly between both states. This separation indicates that Inz-5 binding pushes the Rieske head domain from the *intermediate* to the *c* state. In contrast, the effect of Inz-5 binding on the classification of particle images along *b* state to *intermediate* state component is less distinct (**Fig. 4D**). Previous crystallographic studies have revealed that inhibitor binding in the Q_P_ site can induce small structural changes in the CD2 helices as well as the EF loop, which directly contact the Rieske head domain when it is in the *b* state, and alter the equilibrium of the Rieske head domain (Esser *et al*., 2006; Berry and Huang, 2011). This classification strengthens the above assertion that Inz-5 binds similarly to strobilurin-type inhibitors as it appears to influence the binding pocket, and therefore the Rieske head domain, in a similar manner. Inhibitors in each class often share biological properties such as resistance-conferring mutations (Esser *et al*., 2004), suggesting that cross-resistance to Inz-5 and strobilurin-family inhibitors could occur.

**Fig. 4.**
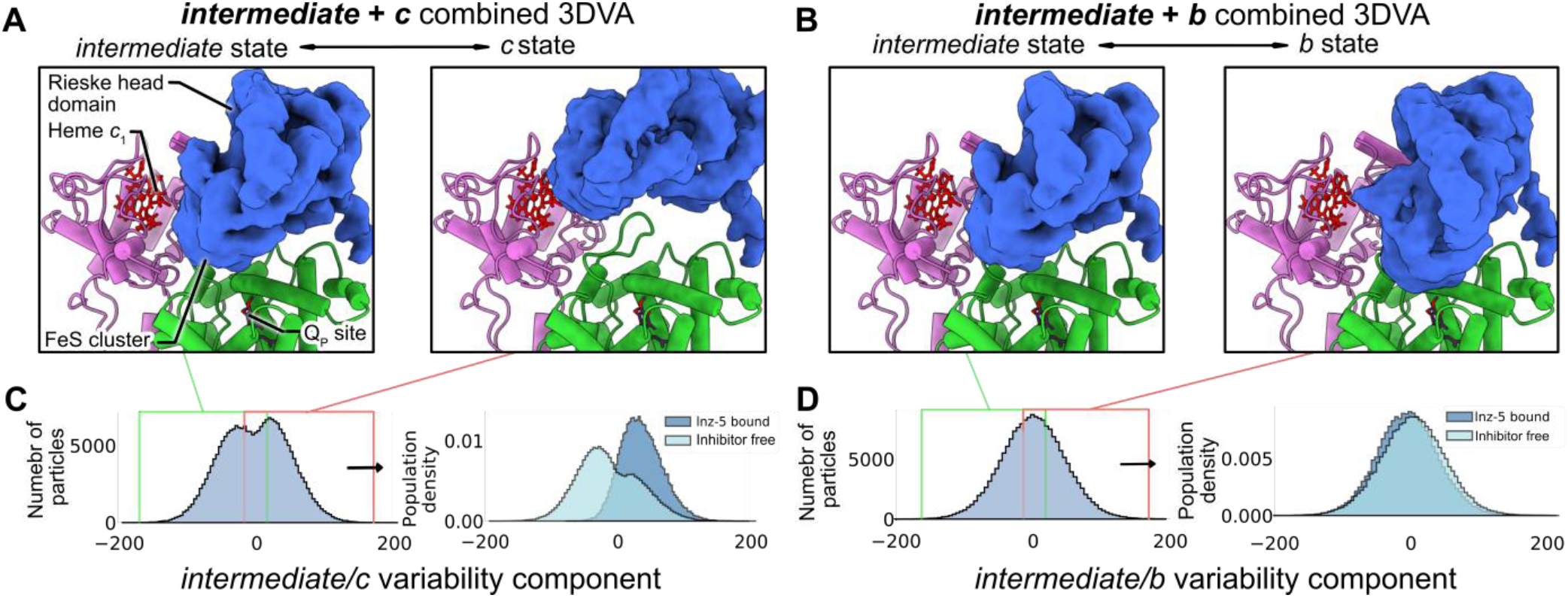
Effect of Inz-5 on Rieske head domain position. The Rieske head domain region of maps calculated from particle images from the extremes of variability components are shown for particle images from the *intermediate* plus *c* population (**A**), and *intermediate* plus *b* population (**B**). Histogram of particle image coefficients from 3DVA of (**C**) *intermediate* plus *c* population and (**D**) *intermediate* plus *b* population. Particle images from the inhibitor-free and Inz-5-bound datasets are shown in the same colour on the left of parts C and D, and different colours in right of parts C and D.

### Rieske protein position is not cooperative between monomers

A major outstanding question regarding the mechanism of CIII_2_ is whether or not the CIII monomers in the dimer function cooperatively. The biphasic binding of stigmatellin to CIII_2_ (Covian *et al*., 2007), along with other arguments based on enzymatic activity, suggested communication between the monomers of CIII_2_ (Covian and Trumpower, 2008). To assess the possibility of coordinated movement of the Rieske head domains across monomers in the dimer, we used 3DVA to sort the conformation of the Rieske head domains in each CIII monomer in the dimer, making use of the two-fold symmetry of the complex. Images were divided into two subsets according to their position along the principal component that indicates whether the Rieske head domain of one monomer is in the *b* or *c* state. For each subset, the position of the Rieske head domain of the opposite monomer was plotted. The results show a lack of correlation between the position of the Rieske head domains across monomers, both for the inhibitor-free (**Fig. 5**) and Inz-5-bound (**Fig. S5**) datasets. This lack of cooperativity, combined with the results above demonstrating the influence of Inz-5 on the Rieske head domain position, infers a lack of cooperativity between Q_P_ sites across the monomers with Inz-5 bound or in the absence of inhibitor. Previous work has suggested that substrate binding to one Q_P_ site causes a decrease in the affinity of the opposite Q_P_ site (Covian *et al*., 2007). It is possible that the cooperativity shown in previous work only occurs for P_f_ inhibitors and not with Inz-5, which is a P_m_ type inhibitor.

**Fig. 5.**
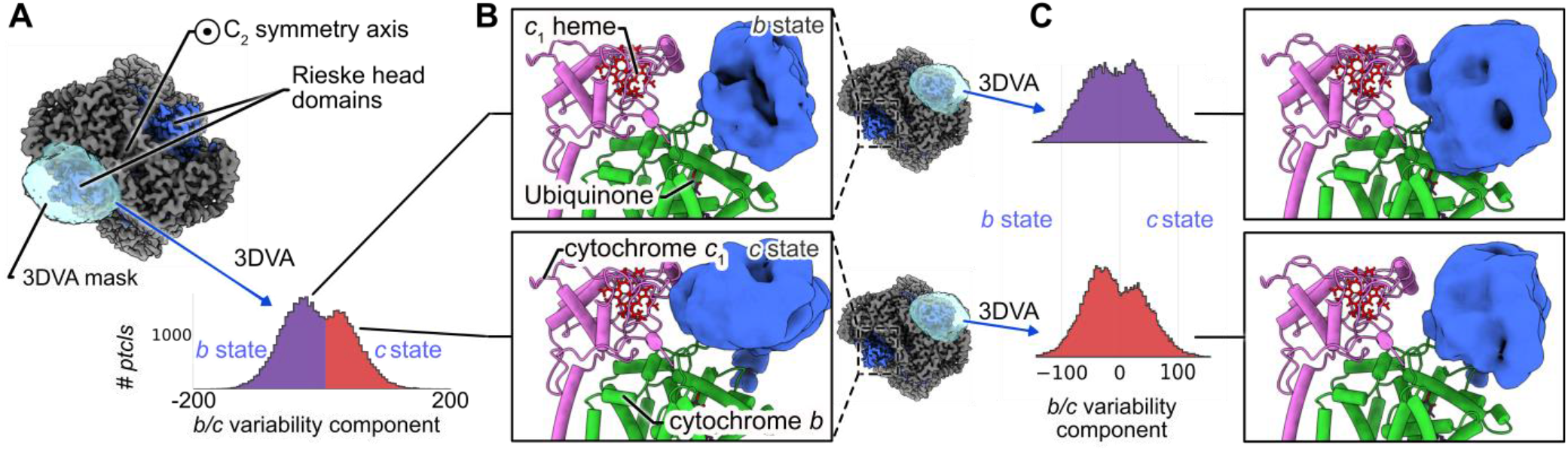
Assessment of cooperativity between CIII_2_ Rieske head domains in opposing monomers for inhibitor free dataset. (**A**) 3DVA of the Rieske head domain from the first monomer was used to divide data into *b* and *c* state (purple and red, respectively). (**B**) Maps of the Rieske head domain region from *b* state (top) and *c* state (bottom) particle images. (**C**) 3DVA of the Rieske head domain of the second monomer from images where first monomer was in the *b* state (top) and *c* state (bottom), with maps produced from all particles from each 3DVA.

## DISCUSSION

The results presented here demonstrate the use of PPCA for the quantitative analysis of conformational equilibria from cryoEM images. Compared to three-dimensional classification and heterogeneous refinement, which has also been used for this purpose (Zhao, Benlekbir and Rubinstein, 2015; Hite and MacKinnon, 2017; Huang *et al*., 2019), PPCA appears to provide a more robust estimate of population differences, possibly because it does not depend on the initial reference maps used. This advantage may also arise because 3DVA accounts for the continuous motion (Punjani and Fleet, 2020) in the Rieske head domain position (Esser *et al*., 2019) rather than attempting to model the motion as a series of discrete states. While a single component effectively models the motion of the Rieske head domain, it should also be possible to analyze movements that can only be described by a combination of components. PPCA may yield components that appear to correspond to coordinated movement of two regions of a protein complex. However, care should be taken when drawing conclusions from these components because there may be other components, which may not be displayed depending on the number of components calculated, that show the same motion except in an anticorrelated fashion.

An earlier atomic model of ovine CI_1_CIII_2_ (Letts *et al*., 2019) also resolved endogenous UQ in the Q_P_ site. While the authors of that study did not comment on the presence of UQ, both the present structure and the earlier structure show UQ in a position similar to stigmatellin. These observations support models of CIII function where UQH_2_ and His161 of the Rieske head domain form a hydrogen bond with UQH_2_ serving as the hydrogen bond donor and His161 as the acceptor (Link and Von Jagow, 1995; Brandt and Okun, 1997; Link, 1997; Crofts, Hong, *et al*., 1999; Hong *et al*., 1999; Rich, 2004). This conformation allows for transfer of the first electron and proton from UQH_2_ to the FeS cluster of the Rieske head domain and His161, respectively. After this electron transfer the Rieske head domain becomes the hydrogen bond donor and semiquinone the acceptor, which holds the Rieske head domain in the *b* position. The interaction is broken only after transfer of the second electron from UQ^•-^ through the *b* chain, allowing the Rieske head domain to travel to the *c* position, reduce cyt. *c*_1_, and release the proton from His161 to the P side of the membrane. In this model, the interaction between the Rieske head domain and UQ^•-^ ensures that the Rieske head domain cannot transfer both electrons from UQH_2_ through the *c* chain (Crofts and Wang, 1989; Jünemann, Heathcote and Rich, 1998; Millett *et al*., 2013; Scharlau *et al*., 2019) and is therefore necessary for CIII to contribute to the PMF.

The results presented here show the presence of three sub-populations of complexes, with the Rieske head domain in the *b, intermediate*, or *c* position. While it is likely that both the Inz-5-bound and drug-free datasets include CIII particles with both occupied and empty Q_P_ sites, the weak UQ density in the drug-free map suggests that Q_P_ sites are mostly empty in that dataset. CryoEM structures of CIII_2_ from other organisms have also found empty Q_P_ sites (Hartley *et al*., 2019; Rathore *et al*., 2019; Maldonado *et al*., 2021; Steimle *et al*., 2021). In an *S. cerevisiae* CIII_2_ structure where the Q_P_ site was empty, the Rieske head domain was observed in an intermediate position (Hartley *et al*., 2019; Rathore *et al*., 2019). Furthermore, recent cryoEM structures of CIII_2_ from *Rhodobacter capsulatus* (Steimle *et al*., 2021) and *Vigna radiata* (Maldonado *et al*., 2021) revealed subpopulations with the FeS domain in either the *b* or *c* positions. Collectively, these data suggest that the FeS head domain stochastically adopts different conformations when the Q_P_ site is unoccupied by ligands (Berry and Huang, 2003; Sarewicz and Osyczka, 2015). Allostery between CIII_2_ dimers has not been reported in any of the above cryoEM studies and it is not detected in the current dataset. However, these results do not preclude the existence of other states in which allosteric interactions may become apparent, either in the presence of specific P_f_ inhibitors (Covian *et al*., 2007), or during catalysis.

The structure of CIII_2_ with Inz-5 bound reveals specific interactions between the inhibitor and residues in the Q_P_ site. Earlier biochemical analysis identified several residues that, when mutated, increased the minimal inhibitory concentration of a similar indazole analogue (Inz-1) (Vincent *et al*., 2016). Replacing cyt. *b* residue Gly143 with Ala, which is in close proximity to the indazole ring of Inz-5, increases the minimum inhibitory concentration for Inz-1 by 5-fold (Vincent *et al*., 2016). This structural alteration also provides protection against the fungicide azoxystrobin without affecting the enzymatic activity of CIII_2_ in *Erysiphe graminis* (Sierotzki, Wullschleger and Gisi, 2000; Bartlett *et al*., 2002). In fact, the specificity of azoxystrobin for the fungal versus plant CIII_2_ has been ascribed to the presence of Ala in the place of Gly143 in the latter (Esser *et al*., 2004). The similarity between the poses of azoxystrobin and Inz-5 binding has important implications for further development of indazole-based antifungals. First, strobilurin derivatives have been reported that possess picomolar affinity for CIII_2_ (Hao *et al*., 2012), suggesting it may be possible to optimize the indazole scaffold to provide similarly potent inhibitors. Second, the classification of both inhibitor classes as P_m_ types, in combination with the large increase in the minimal inhibitory concentration observed in the Gly143Ala mutation, suggest that novel indazole derivatives may be useful as agricultural fungicides in addition to the medicinal applications originally envisioned for them.

In *C. albicans*, the human pathogen most often responsible for life-threatening fungal infections, mutation of Leu275 to Phe in the Q_P_ site of cyt. *b* leads to a large increase in the minimal inhibitory concentration of Inz-1 (Vincent *et al*., 2016). The structure presented here explains this finding as it shows that the indazole moiety of Inz-5 is in close contact with Leu275. Interestingly, mammalian and *R. sphaeroides* CIII_2_ contain Phe at this position, but crystal structures of the bovine and *R. sphaeroides* CIII_2_ in complex with azoxystrobin or MOAS show that binding of these inhibitors is not disrupted. Therefore, the increase in minimum inhibitory concentration for Inz-1 in the Leu275Phe mutant may be due to the position of the indazole’s fused ring near Leu275, which is larger than the single ring α-benzene and phenyl group of MOAS and azoxystrobin, respectively. Importantly, these data indicate that the presence of a Phe at position 275 in the bovine and human enzymes, as compared to Leu in *C. albicans*, explains the fungal selectivity of indazole derivatives (Vincent *et al*., 2016). Infections by fungi pose an under-appreciated but escalating health problem, with a global disease burden now affecting billions and killing more than 1.5 million people annually (Fisher *et al*., 2018). The structural insights provided here highlight promising avenues to improve the antifungal potency and selectivity of indazole-based inhibitors of CIII and can ultimately help generate new, more effective antifungals.

## Supporting information

Rieske-HD dynamics

Inz-5 binding pose

## ACCESSION NUMBERS

All cryoEM maps described in this article have been deposited in the Electron Microscopy Data Bank (EMD-XXXX to EMD-XXXX). Atomic models have been deposited to the PDB (PDB IDs: XXXX and XXXX)

## ACKNOWLEDGEMENTS

JDT was supported by a postdoctoral fellowship from the Canadian Institutes of Health Research (CIHR), ZL was supported by a Precision Medicine Initiative (PRiME) internal fellowship (PRMF2020-005), JLR and LEC were supported by the Canada Research Chairs program, and LEC was supported by the Canadian Institutes for Advanced Research. This research was supported by CIHR Grant PJT162186 (JLR), CIHR Grant FDN154288 (LEC), the Knut and Alice Wallenberg Foundation grant 2019.0043 (PB), and the Swedish Research Council grant 2018-04619 (PB). Cryo-EM data was collected at the Toronto High-Resolution High-Throughput cryo-EM facility, supported by the Canada Foundation for Innovation and Ontario Research Fund.

## AUTHOR CONTRIBUTIONS

JDT assisted with CIII_2_ purification, prepared EM grids, collected and analysed EM data, and analysed enzyme inhibition data, ZL prepared CIII_2_ strains, purified CIII_2_, and performed enzyme inhibition experiments. JLR, LEC, PB, and LW supervised the research. JDT and JLR wrote the manuscript and prepared the figures with input from the other authors.

## STAR METHODS

### Strain and cell growth

The *Candida albicans* strain CaLC5421, with both alleles of the *QCR2* gene modified to encode a C-terminal HIS_6_-3×FLAG, was constructed by homologous recombination. Briefly, PCR products amplified from oLC6938 (5’ GAAGAAGCCCCAATTTCTAAATTCAACTATGTTGCTGTTGGTGATCTTGATGTGTTACCATATGCTGATGAATTGggtcgacggatccc c 3’) and oLC6939 (5’ AATAAATCAGTAATCTATTACGTACATGAATGTTTATTTTATTCACATATGACTTAAATAAACTGCTAAAGGGtcgatgaattcgagctc g 3’) from pLC1084 (pFA-HIS_6_-3×FLAG -*SAT1* (Zhang *et al*., 2012)) and pLC1086 (pFA-HIS_6_-3×FLAG-*HIS1* (Zhang *et al*., 2012)) were introduced sequentially into a laboratory reference strain of *C. albicans* (SN95). CaLC5421 was cultured in YPGalactose (2% [w/v] peptone, 1% [w/v] yeast extract, 2% [w/v] galactose supplemented with 0.1 mM uridine) to mid log phase while shaking at 200 rpm at 30°C before harvested by centrifugation at 3000 ×g (5 min, 4 °C).

### Preparation of mitochondrial membranes and Isolation of CIII_2_

Crude mitochondria were prepared from CaLC5421 as described (Meisinger, Pfanner and Truscott, 2006). After addition of 10 mM DTT, followed by zymolyase (Bioshop; 3 mg per gram wet weight) at 30 °C, cells were lysed by homogenization in ice cold hypotonic buffer containing 0.6 M sorbitol, 10 mM Tris-HCl (pH 7.4), 1 mM EDTA, 1 mM PMSF, and 0.2% [w/v] BSA (A7030 (Sigma-Aldrich)). Mitochondria were pelleted by centrifugation at 12,000 ×g (25 min, 4 °C) and stored at -80 °C in SEM buffer (10 mM MOPS-KOH pH 7.2, 250 mM sucrose, 1 mM EDTA, 1 mM PMSF) at a total protein concentration of ∼10 mg/mL (determined by Bio-Rad DC protein assays after solubilization in 1% [w/v] SDS). To prepare mitochondrial membranes, mitochondria were homogenized with a Dounce homogenizer in solubilization buffer (SB; 50 mM potassium phosphate [KPi] pH 7.4, 100 mM KCl, 5 mM 6-aminocaproic acid, 5 mM 4-aminobenzamidine, 1 mM PMSF). Membranes were pelleted by centrifugation at 48,000 ×g (45 min, 4 °C) and resuspended in SB to a total protein concentration of ∼10 mg/mL. Membranes were then solubilized by addition of n-dodecyl β-D-maltoside (DDM (Bioshop); 0.8 mg /mg protein) and incubated for 1 h with gentle stirring at 4°C. Insoluble materials were removed by centrifugation at 180,000 ×g for 45min. The cleared supernatant was diluted with SB supplemented with 0.02% [w/v] DDM to reduce DDM concentration to ∼0.2% [w/v] before incubated with 1 mL ANTI-FLAG M2 Affinity Gel (Sigma) for 2 h at 4 °C. The affinity resin with protein bound was washed with 30 bed volumes of wash buffer (KPi pH 7.4, 100 mM KCl, 0.02% [w/v] DDM). CIII_2_ was eluted with the wash buffer supplemented with 150 µg/mL 3×FLAG peptide (APExBIO) and concentrated on Vivaspin 6 (100 kDa MWCO; GE Healthcare). Buffer was exchanged to 20 mM KPi pH 7.4 containing 100 mM KCl and 0.01% [w/v] glyco-diosgenin (GDN; Avanti) by two rounds of dilution (1:20) and concentration on the same type of concentrator. CIII_2_ was purified further by size exclusion chromatography with an Äkta Pure (GE Healthcare) operated at 4 °C with UV detection at 280 nm and 420 nm. The sample was loaded on a Superose 6 Increase 10/300 GL column (GE Healthcare) equilibrated with 100 mM KCl, 20 mM KPi buffer, pH 7.4, 0.01% [w/v] GDN. Fractions containing CIII_2_ were collected and concentrated to ∼4.5 mg/ml (9 µM) studies of the inhibitor-free complex, and ∼8 mg/ml (16 µM) for studies of the Inz-5-bound specimen. UV-Vis spectra were recorded with a NanoDrop 1000 (Thermo Scientific) immediately before grid preparation. To bind Inz-5, Inz-5 dissolved in DMSO was diluted by via serial dilution in buffer and then added to the CIII_2_ sample at a final concentration of 29 µM (0.07% [v/v] DMSO).

### Grid preparation and cryoEM

Purified CIII_2_ (2 µl at ∼4.5 mg/ml for inhibitor-free complex, ∼8 mg/ml for Inz-5 bound) was applied to homemade nanofabricated holey gold grids (Marr, Benlekbir and Rubinstein, 2014; Russo and Passmore, 2014) that had previously been glow-discharged in air (120 s, 20 mA with a PELCO easiGlow). Grids were blotted for 28 s at 4 °C and 100% humidity before freezing in a liquid propane/ethane mixture with a modified Vitrobot mark III (FEI). CryoEM data were collected with a Titan Krios G3 electron microscope (Thermo Fisher Scientific) operated at 300 kV and equipped with a prototype Falcon 4 direct detector device camera. Automated data collection was done with the EPU software package. A dataset of 3,953 movies was collected for the inhibitor-free sample and a dataset of 4,396 movies was collected for the Inz-5-bound sample, with each movie consisting of 29 exposure fractions over 8.7 s. The nominal magnification was 75,000× with a calibrated pixel size of 1.03 Å and the camera exposure rate and the total exposure were 4.6 e^-^/pixel/s and ∼42 e^-^/Å^2^, respectively (**Table S1**).

### CryoEM image processing

All image analysis was performed within cryoSPARC ver. 2 (Punjani *et al*., 2017). Movies were aligned with *MotionCor2* (Zheng *et al*., 2017) and contrast transfer function (CTF) parameters were estimated in patches with a 7×7 grid. The dataset was manually curated to remove movies with devitrified ice, large cracks in the ice, or poor CTF fit parameters, reducing the dataset size to 3,634 and 3,652 movies for the inhibitor-free and Inz-5-bound datasets, respectively. Templates for particle selection were generated by 2D classification of manually-selected particle images and reference-based particle selection led to 1,151,426 and 1,129,413 particle images for the inhibitor-free and Inz-5-bound datasets, respectively. Particle images were corrected for individual particle motion and extracted in 288×288 pixel boxes (**Table S1**). Extracted particle images were cleaned with 6 to 8 rounds of *ab initio* 3D classification and heterogeneous refinement, keeping only the particle images that gave class averages corresponding to CIII_2_ after each round. This procedure further reduced the size of the datasets to 96,783 and 61,067 particle images for the inhibitor-free and Inz-5-bound datasets, respectively. Final maps were generated with non-uniform refinement enforcing C_2_ symmetry. These maps were analyzed with 3D variability analysis (3DVA) and cluster analysis (Punjani and Fleet, 2020) within a mask selecting the region of the map corresponding to Rieske head domain. The first round of 3DVA and cluster analysis led to two populations: the first with the Rieske head domain in the *c* state, and the second a mix of the *b* and *intermediate*-state particle images. A second round of 3DVA on the mixed *b/intermediate*-state particle images allowed for separation into two populations. This analysis was performed for both the inhibitor-free and Inz-5-bound datasets (See **Fig. S3**).

### Atomic model building

Homology models were created with the web-based SWISS-MODEL service (Waterhouse *et al*., 2018) starting with a model of CIII_2_ from *S. cerevisiae* (Hunte *et al*., 2000). Refinement of the homology model was performed in *coot* (Emsley *et al*., 2010), *ISOLDE* (Croll, 2018), and *phenix* (Liebschner *et al*., 2019). Final refinement and calculation of atomic displacement parameters were done with real space refine in *phenix* (for model building summary see **Table S2**). Movies were created using UCSF ChimeraX and HitFilm Express.

### CIII_2_ activity assays and IC_50_ calculation

CIII_2_ activity was measured by following the reduction of equine cytochrome *c* (Sigma) spectrophotometrically using decylubiquinol (DBH_2_) as the electron donor. DBH_2_ was generated from decylubiquinone (Sigma) as described (Trumpower and Edwards, 1979; Covian, Gutierrez-Cirlos and Trumpower, 2004). A reaction mixture (50 µL/per well) containing 150 µM cytochrome *c* and ∼25 nM purified CIII_2_ in buffer (50 mM KPi pH 7.4, 100 mM KCl, 0.1 mM EDTA, 0.5 mM KCN, 0.01% [w/v] GDN) was deposited into the wells of 96-well plate. 2.5 mM Inz-5 stock in DMSO was added to the required concentration with a D300e digital dispenser (Tecan). A compensatory amount of DMSO was added to each well to ensure they all had the same amount (0.033% [v/v]). After incubation at room temperature for 5 min, reactions were initiated by addition of 100 µl of ∼120 µM DBH_2_ in ethanol into the reaction buffer. Absorbance at 550 nm was recorded every 15 seconds with SpectraMax plate reader (Molecular Devices). The IC_50_ was calculated with a custom Python script and errors were estimated using Monte Carlo simulations (Koehler, Brown and Haneuse, 2009). Briefly, the *RMSD* between data and the best fit curve was calculated and simulated data sets were created using the best fit parameters. Random errors *N*(0, *RMSD*) (normal random numbers with standard deviation = *RMSD*) were added to each data point and the simulated data with errors was fit to extract an IC_50_. The addition of random errors and fitting was repeated 10,000 times and the standard deviation of the 10,000 best fit parameters was taken to be the standard deviation of the IC_50_.

### Assessment of the Rieske head domain equilibrium changes associated with Inz-5 binding

To assess the effect of Inz-5 on the Rieske head domain position equilibrium, the inhibitor-free and Inz-5-bound datasets were combined and subject to non-uniform refinement while enforcing C_2_ symmetry, followed by symmetry expansion and 3DVA in order to cluster the particles into *c*-, *b*-, and *intermediate*-state populations (**Fig. S3**). The *b/intermediate*-state particle images, as well as the *c/intermediate*-state particle images, were combined and subject to 3DVA. The 3DVA results were analyzed with custom Python scripts that sorted particle images according to which dataset they originated from and plotted the image coefficients for each particle image along the first vector. The intermediate output for each 3DVA job was used to generate a map of the Rieske head domain in order to ensure that the PPCA correctly captured the motion between the two states of interest (the *b* state to the *intermediate* state, and the *c* state to the *intermediate* state). The image coefficient for each dataset were plotted in different colors along the vector in order to assess the change in Rieske head domain equilibrium upon Inz-5 binding.

### Assessment of cooperativity between Rieske head domains across monomers

Particle images from the combined dataset were split into the three populations (*c, b*, and *intermediate* Rieske head domain positions) using the two rounds of 3DVA processing described above (**Fig. S3**) and then analyzed with a custom python script. First, the particle images were divided into two datasets (inhibitor free and Inz-5 bound), which were analyzed separately. Next, all particle images with the Rieske head domain in the *intermediate* state were discarded. Remaining particle images were analysed by 3DVA, yielding a *c* to *b* state vector, and two image coefficients for every particle image, as each CIII_2_ particle image contained information for two Rieske head domains. The image coefficients for each particle image were sorted, at random, into two groups so that each group contained information for one CIII_2_ monomer Rieske head domain. The image coefficients for the first monomer were plotted along the first vector, and the particle images were clustered according to these image coefficients. Local refinement of the first monomer for each cluster separately confirmed the clusters contained *b* state and *c* state particles. The image coefficients for the second monomer were plotted as two histograms, one for the particles images that contained a Rieske head domain from the first monomer in the *b* state and another for the *c* state. Local refinement was also performed for the second monomer from each cluster to confirm the position of the Rieske head domain.

## REFERENCES

Andries, K. et al. (2005) ‘A Diarylquinoline Drug Active on the ATP Synthase of Mycobacterium tuberculosis’, Science, 307(5707), pp. 223 LP–227. doi: 10.1126/science.1106753.

Bartlett, D. W. et al. (2002) ‘The strobilurin fungicides’, Pest Management Science, 58(7), pp. 649–662. doi: 10.1002/ps.520.

Berry, E. A. and Huang, L. S. (2003) ‘Observations concerning the quinol oxidation site of the cytochrome bc 1 complex’, FEBS Letters, 555(1), pp. 13–20. doi: 10.1016/S0014-5793(03)01099-8.

Berry, E. A. and Huang, L. S. (2011) ‘Conformationally linked interaction in the cytochrome bc1 complex between inhibitors of the Qo site and the Rieske iron-sulfur protein’, Biochimica et Biophysica Acta - Bioenergetics. Elsevier B.V., 1807(10), pp. 1349–1363. doi: 10.1016/j.bbabio.2011.04.005.

Brandt, U. and von Jagow, G. (1991) ‘Analysis of inhibitor binding to the mitochondrial cytochrome c reductase by fluorescence quench titration: Evidence for a “catalytic switch” at the Qo center’, European Journal of Biochemistry, 195(1), pp. 163–170. doi: 10.1111/j.1432-1033.1991.tb15690.x.

Brandt, U. and Okun, J. G. (1997) ‘Role of deprotonation events in ubihydroquinone:cytochrome c oxidoreductase from bovine heart and yeast mitochondria’, Biochemistry, 36(37), pp. 11234–11240. doi: 10.1021/bi970968g.

Burley, A. S. K. and Petsko, G. a (1985) ‘Aromatic-Aromatic Interaction: A Mechanism of Protein Structure Stabilization’, Science, 229(4708), pp. 23–28.

Covian, R. et al. (2007) ‘Asymmetric binding of stigmatellin to the dimeric Paracoccus denitrificans bc1 complex: Evidence for anti-cooperative ubiquinol oxidation and communication between center P ubiquinol oxidation sites’, Journal of Biological Chemistry, 282(31), pp. 22289–22297. doi: 10.1074/jbc.M702132200.

Covian, R., Gutierrez-Cirlos, E. B. and Trumpower, B. L. (2004) ‘Anti-cooperative Oxidation of Ubiquinol by the Yeast Cytochrome bc 1 Complex’, Journal of Biological Chemistry, 279(15), pp. 15040–15049. doi: 10.1074/jbc.M400193200.

Covian, R. and Trumpower, B. L. (2008) ‘Biochimica et Biophysica Acta Regulatory interactions in the dimeric cytochrome bc 1 complex : The advantages of being a twin’, 1777, pp. 1079–1091. doi: 10.1016/j.bbabio.2008.04.022.

Crofts, A. R., Guergova-Kuras, M., et al. (1999) ‘Mechanism of ubiquinol oxidation by the bc1 complex: Role of the iron sulfur protein and its mobility’, Biochemistry, 38(48), pp. 15791–15806. doi: 10.1021/bi990961u.

Crofts, A. R., Hong, S., et al. (1999) ‘Pathways for proton release during ubihydroquinone oxidation by the bc1 complex’, Proceedings of the National Academy of Sciences of the United States of America, 96(18), pp. 10021–10026. doi: 10.1073/pnas.96.18.10021.

Crofts, A. R. and Wang, Z. (1989) ‘How rapid are the internal reactions of the ubiquinol:cytochrome c2 oxidoreductase?’, Photosynthesis Research, 22(1), pp. 69–87. doi: 10.1007/BF00114768.

Croll, T. I. (2018) ‘ISOLDE: a physically realistic environment for model building into low-resolution electron-density maps’, Acta Crystallographica Section D, 74(6), pp. 519–530. doi: 10.1107/S2059798318002425.

Darrouzet, E. et al. (2001) ‘Large scale domain movement in cytochrome bc1: A new device for electron transfer in proteins’, Trends in Biochemical Sciences, 26(7), pp. 445–451. doi: 10.1016/S0968-0004(01)01897-7.

Emsley, P. et al. (2010) ‘Features and development of Coot’, Acta Crystallographica Section D: Biological Crystallography. International Union of Crystallography, 66(4), pp. 486–501. doi: 10.1107/S0907444910007493.

Esser, L. et al. (2004) ‘Crystallographic studies of quinol oxidation site inhibitors: A modified classification of inhibitors for the cytochrome bc1 complex’, Journal of Molecular Biology, 341(1), pp. 281–302. doi: 10.1016/j.jmb.2004.05.065.

Esser, L. et al. (2006) ‘Surface-modulated motion switch: Capture and release of iron-sulfur protein in the cytochrome bc1 complex’, Proceedings of the National Academy of Sciences of the United States of America, 103(35), pp. 13045–13050. doi: 10.1073/pnas.0601149103.

Esser, L. et al. (2019) ‘Crystal structure of bacterial cytochrome bc1 in complex with azoxystrobin reveals a conformational switch of the Rieske iron–sulfur protein subunit’, Journal of Biological Chemistry, 294(32), pp. 12007–12019. doi: 10.1074/jbc.RA119.008381.

Fisher, M. C. et al. (2018) ‘Worldwide emergence of resistance to antifungal drugs challenges human health and food security’, Science, 360(6390), pp. 739 LP–742. doi: 10.1126/science.aap7999.

Hao, G. F. et al. (2012) ‘Computational discovery of picomolar Q o site inhibitors of cytochrome bc 1 complex’, Journal of the American Chemical Society, 134(27), pp. 11168–11176. doi: 10.1021/ja3001908.

Hartley, A. M. et al. (2019) ‘Structure of yeast cytochrome c oxidase in a supercomplex with cytochrome bc 1’, Nature Structural and Molecular Biology, 26(1), pp. 78–83. doi: 10.1038/s41594-018-0172-z.

Hite, R. K. and MacKinnon, R. (2017) ‘Structural Titration of Slo2.2, a Na+-Dependent K+ Channel’, Cell. Elsevier, 168(3), pp. 390-399.e11. doi: 10.1016/j.cell.2016.12.030.

Hong, S. et al. (1999) ‘The energy landscape for ubihydroquinone oxidation at the Q(o) site of the bc1 complex in Rhodobacter sphaeroides’, Journal of Biological Chemistry, 274(48), pp. 33931–33944. doi: 10.1074/jbc.274.48.33931.

Huang, L.-S. et al. (2005) ‘Binding of the respiratory chain inhibitor antimycin to the mitochondrial bc1 complex: a new crystal structure reveals an altered intramolecular hydrogen-bonding pattern’, Journal of molecular biology, 351(3), pp. 573–597. doi: 10.1016/j.jmb.2005.05.053.

Huang, R. et al. (2016) ‘Unfolding the mechanism of the AAA+ unfoldase VAT by a combined cryo-EM, solution NMR study’, Proceedings of the National Academy of Sciences of the United States of America, 113(29), pp. E4090–W4199. doi: 10.1073/pnas.1603980113.

Huang, R. et al. (2019) ‘Cooperative subunit dynamics modulate p97 function’, Proceedings of the National Academy of Sciences of the United States of America, 116(1), pp. 158–167. doi: 10.1073/pnas.1815495116.

Hunte, C. et al. (2000) ‘Structure at 2.3 Å resolution of the cytochrome bc1 complex from the yeast Saccharomyces cerevisiae co-crystallized with an antibody Fv fragment’, Structure, 8(6), pp. 669–684. doi: 10.1016/S0969-2126(00)00152-0.

Iwata, S. et al. (1998) ‘Complete structure of the 11-subunit bovine mitochondrial cytochrome bc1 complex’, Science, 281(5373), pp. 64–71. doi: 10.1126/science.281.5373.64.

Jünemann, S., Heathcote, P. and Rich, P. R. (1998) ‘On the mechanism of quinol oxidation in the bc1 complex’, Journal of Biological Chemistry, 273(34), pp. 21603–21607. doi: 10.1074/jbc.273.34.21603.

Kim, H. et al. (1998) ‘Inhibitor binding changes domain mobility in the iron-sulfur protein of the mitochondrial bc1 complex from bovine heart’, Proceedings of the National Academy of Sciences of the United States of America, 95(14), pp. 8026–8033. doi: 10.1073/pnas.95.14.8026.

Koehler, E., Brown, E. and Haneuse, S. J. P. A. (2009) ‘On the assessment of Monte Carlo error in simulation-based Statistical analyses’, American Statistician, 63(2), pp. 155–162. doi: 10.1198/tast.2009.0030.

Lange, C. and Hunte, C. (2002) ‘Crystal structure of the yeast cytochrome bc1 complex with its bound substrate cytochrome c’, Proceedings of the National Academy of Sciences of the United States of America, 99(5), pp. 2800–2805. doi: 10.1073/pnas.052704699.

Letts, J. A. et al. (2019) ‘Structures of Respiratory Supercomplex I+III2 Reveal Functional and Conformational Crosstalk’, Molecular Cell. Elsevier Inc., 75(6), pp. 1131-1146.e6. doi: 10.1016/j.molcel.2019.07.022.

Liebschner, D. et al. (2019) ‘Macromolecular structure determination using X-rays, neutrons and electrons: recent developments in Phenix’, Acta Crystallographica Section D, 75(10), pp. 861–877. doi: 10.1107/S2059798319011471.

Link, T. A. (1997) ‘The role of the “Rieske” iron sulfur protein in the hydroquinone oxidation (Qp) site of the cytochrome bc1 complex’, FEBS Letters. Federation of European Biochemical Societies, 412(2), pp. 257–264. doi: 10.1016/S0014-5793(97)00772-2.

Link, T. A. and Iwata, S. (1996) ‘Functional implications of the structure of the “Rieske” iron-sulfur protein of bovine heart mitochondrial cytochrome bc1 complex’, Biochimica et Biophysica Acta - Bioenergetics, 1275(1–2), pp. 54–60. doi: 10.1016/0005-2728(96)00050-3.

Link, T. A. and Von Jagow, G. (1995) ‘Zinc ions inhibit the QP center of bovine heart mitochondrial bc1 complex by blocking a protonatable group’, Journal of Biological Chemistry, 270(42), pp. 25001–25006. doi: 10.1074/jbc.270.42.25001.

Maldonado, M. et al. (2021) ‘Atomic structures of respiratory complex III2, complex IV, and supercomplex III2-IV from vascular plants’, eLife, 10, pp. 1–34. doi: 10.7554/eLife.62047.

Marr, C. R., Benlekbir, S. and Rubinstein, J. L. (2014) ‘Fabrication of carbon films with ∼500nm holes for cryo-EM with a direct detector device’, Journal of Structural Biology. Elsevier Inc., 185(1), pp. 42–47. doi: 10.1016/j.jsb.2013.11.002.

Mazu, T. K. et al. (2016) ‘The Mechanistic Targets of Antifungal Agents: An Overview’, Mini reviews in medicinal chemistry, 16(7), pp. 555–578. doi: 10.2174/1389557516666160118112103.

Meisinger, C., Pfanner, N. and Truscott, K. N. (2006) ‘Isolation of yeast mitochondria.’, Methods in molecular biology (Clifton, N.J.), 313(1), pp. 33–39. doi: 10.1385/1-59259-958-3:033.

Millett, F. et al. (2013) ‘Design and use of photoactive ruthenium complexes to study electron transfer within cytochrome bc1 and from cytochrome bc1 to cytochrome c’, Biochimica et Biophysica Acta - Bioenergetics. Elsevier B.V., 1827(11–12), pp. 1309–1319. doi: 10.1016/j.bbabio.2012.09.002.

Osyczka, A., Moser, C. C. and Dutton, P. L. (2005) ‘Fixing the Q cycle’, Trends in Biochemical Sciences, 30(4), pp. 176–182. doi: 10.1016/j.tibs.2005.02.001.

Pfaller, M. A. and Diekema, D. J. (2010) Epidemiology of invasive mycoses in North America, Critical Reviews in Microbiology. doi: 10.3109/10408410903241444.

Punjani, A. et al. (2017) ‘CryoSPARC: Algorithms for rapid unsupervised cryo-EM structure determination’, Nature Methods, 14(3), pp. 290–296. doi: 10.1038/nmeth.4169.

Punjani, A. and Fleet, D. J. (2020) ‘3D Variability Analysis: Directly resolving continuous flexibility and discrete heterogeneity from single particle cryo-EM images’, bioRxiv, p. 2020.04.08.032466. doi: 10.1101/2020.04.08.032466.

Rathore, S. et al. (2019) ‘Cryo-EM structure of the yeast respiratory supercomplex’, Nature Structural and Molecular Biology. Springer US, 26(1), pp. 50–57. doi: 10.1038/s41594-018-0169-7.

Rich, P. R. (2004) ‘The quinone chemistry of bc complexes’, Biochimica et Biophysica Acta - Bioenergetics, 1658(1–2), pp. 165–171. doi: 10.1016/j.bbabio.2004.04.021.

Roemer, T. and Krysan, D. J. (2014) ‘Antifungal Drug Development: Challenges, Unmet Clinical Needs, and New Approaches’, Cold Spring Harb Perspect Med, 4, p. a019703.

Russo, C. J. and Passmore, L. A. (2014) ‘Ultrastable gold substrates for electron cryomicroscopy’, Science, 346(6215), pp. 1377 LP–1380. doi: 10.1126/science.1259530.

Sarewicz, M. and Osyczka, A. (2015) ‘Electronic connection between the quinone and cytochrome c redox pools and its role in regulation of mitochondrial electron transport and redox signaling’, Physiological Reviews, 95(1), pp. 219–243. doi: 10.1152/physrev.00006.2014.

Scharlau, M. et al. (2019) ‘Definition of the Interaction Domain and Electron Transfer Route between Cytochrome c and Cytochrome Oxidase’, Biochemistry. American Chemical Society, 58(40), pp. 4125–4135. doi: 10.1021/acs.biochem.9b00646.

Sierotzki, H., Wullschleger, J. and Gisi, U. (2000) ‘Point mutation in cytochrome b gene conferring resistance to strobilurin fungicides in Erysiphe graminis f. Sp. Tritici field isolates’, Pesticide Biochemistry and Physiology, 68(2), pp. 107–112. doi: 10.1006/pest.2000.2506.

Steimle, S. et al. (2021) ‘Cryo-EM structures of engineered active bc1-cbb3 type CIII2CIV super-complexes and electronic communication between the complexes’, Nature Communications. Springer US, 12(1). doi: 10.1038/s41467-021-21051-4.

Thierbach, G. and Reichenbach, H. (1981) ‘Myxothiazol, a new inhibitor of the cytochrome b-c1 segment of the respiratory chain’, BBA - Bioenergetics, 638(2), pp. 282–289. doi: 10.1016/0005-2728(81)90238-3.

Tian, H. et al. (1999) ‘Evidence for the head domain movement of the Rieske iron-sulfur protein in electron transfer reaction of the cytochrome bc1 complex’, Journal of Biological Chemistry, 274(11), pp. 7146–7152. doi: 10.1074/jbc.274.11.7146.

Trumpower, B. L. and Edwards, C. A. (1979) ‘Purification of a reconstitutively active iron-sulfur protein (oxidation factor) from succinate. cytochrome c reductase complex of bovine heart mitochondria.’, Journal of Biological Chemistry, 254(17), pp. 8697–8706. doi: 10.1016/S0021-9258(19)86947-8.

Vincent, B. M. et al. (2016) ‘A Fungal-Selective Cytochrome bc1 Inhibitor Impairs Virulence and Prevents the Evolution of Drug Resistance’, Cell Chemical Biology. Elsevier Ltd, 23(8), pp. 978–991. doi: 10.1016/j.chembiol.2016.06.016.

Waterhouse, A. et al. (2018) ‘SWISS-MODEL: Homology modelling of protein structures and complexes’, Nucleic Acids Research. Oxford University Press, 46(W1), pp. W296–W303. doi: 10.1093/nar/gky427.

Wikström, M. K. F. and Berden, J. A. (1972) ‘Oxidoreduction of cytochrome b in the presence of antimycin’, Biochimica et Biophysica Acta (BBA) - Bioenergetics, 283(3), pp. 403–420. doi: https://doi.org/10.1016/0005-2728(72)90258-7.

Xia, D. et al. (2013) ‘Structural analysis of cytochrome bc1 complexes: Implications to the mechanism of function’, Biochimica et Biophysica Acta - Bioenergetics. Elsevier B.V., pp. 1278–1294. doi: 10.1016/j.bbabio.2012.11.008.

Xia, K., Yu, L. and Yu, C. A. (2000) ‘Confirmation of the involvement of protein domain movement during the catalytic cycle of the cytochrome bc1 complex by the formation of an intersubunit disulfide bond between cytochrome b and the iron-sulfur protein’, Journal of Biological Chemistry, 275(49), pp. 38597–38604. doi: 10.1074/jbc.M007444200.

Zhang, A. et al. (2012) ‘The Tlo proteins are stoichiometric components of Candida albicans Mediator anchored via the Med3 subunit’, Eukaryotic Cell, 11(7), pp. 874–884. doi: 10.1128/EC.00095-12.

Zhang, C. et al. (2020) ‘Ecotoxicology of strobilurin fungicides’, Science of the Total Environment. Elsevier B.V., 742, p. 140611. doi: 10.1016/j.scitotenv.2020.140611.

Zhang, Z. et al. (1998) ‘Electron transfer by domain movement in cytochrome bc1’, Nature, 392(6677), pp. 677–684. doi: 10.1038/33612.

Zhao, J., Benlekbir, S. and Rubinstein, J. L. (2015) ‘Electron cryomicroscopy observation of rotational states in a eukaryotic V-ATPase’, Nature, 521(7551), pp. 241–245. doi: 10.1038/nature14365.

Zheng, S. Q. et al. (2017) ‘MotionCor2: Anisotropic correction of beam-induced motion for improved cryo-electron microscopy’, Nature Methods. Nature Publishing Group, 14(4), pp. 331–332. doi: 10.1038/nmeth.4193.

