## Supplemental figures for "Rieske head domain dynamics and indazole-derivative inhibition of *Candida albicans* complex III"

**Table S1. Cryo-EM data acquisition and image processing.**

| | |
|---|---|
| **Data Collection** | |
| Electron Microscope | Titan Krios |
| Camera | Falcon 4 |
| Voltage (kV) | 300 |
| Nominal Magnification | 75,000 |
| Calibrated physical pixel size (Å) | 1.03 |
| Total exposure (e/Å$^2$) | 42 |
| Exposure rate (e/pixel/s) | 4.6 |
| Number of frames | 29 |
| Defocus range (µm) | 0.9 to 2 |
| **Image Processing** | |
| Motion correction software | *MotionCor2* |
| CTF estimation software | *cryoSPARC v2* |
| Particle selection software | *cryoSPARC v2* |
| Micrographs used in inhibitor free dataset | 3,953 |
| Micrographs used in Inz-5 bound dataset | 4,396 |
| Particle images selected in inhibitor free dataset | 1,151,426 |
| Particle images selected in Inz-5 bound dataset | 1,129,413 |
| 3D map classification and refinement software | *cryoSPARC v2* |

**Table S2. CryoEM map and atomic model statistics.**

| Dataset | Inhibitor free | Inz-5 bound |
|---|---|---|
| Associated PDB ID | | |
| Modelling and refinement software | Coot, phenix, ISOLDE | Coot, phenix, ISOLDE |
| Protein residues | 1906 | 1909 |
| Ligand | HEM:3, FES:1, UQ:2 | HEM:3, FES:1, INZ:1 |
| RMSD bond length (Å) | 0.004 | 0.003 |
| RMSD bond angle (°) | 0.677 | 0.691 |
| Ramachandran outliers (%) | 0 | 0 |
| Ramachandran favoured (%) | 96.13 | 97.25 |
| Rotamer outliers (%) | 0 | 0 |
| Clash score | 8.92 | 8.97 |
| MolProbability score | 1.74 | 1.62 |
| EMringer score | 4.25 | 4.04 |

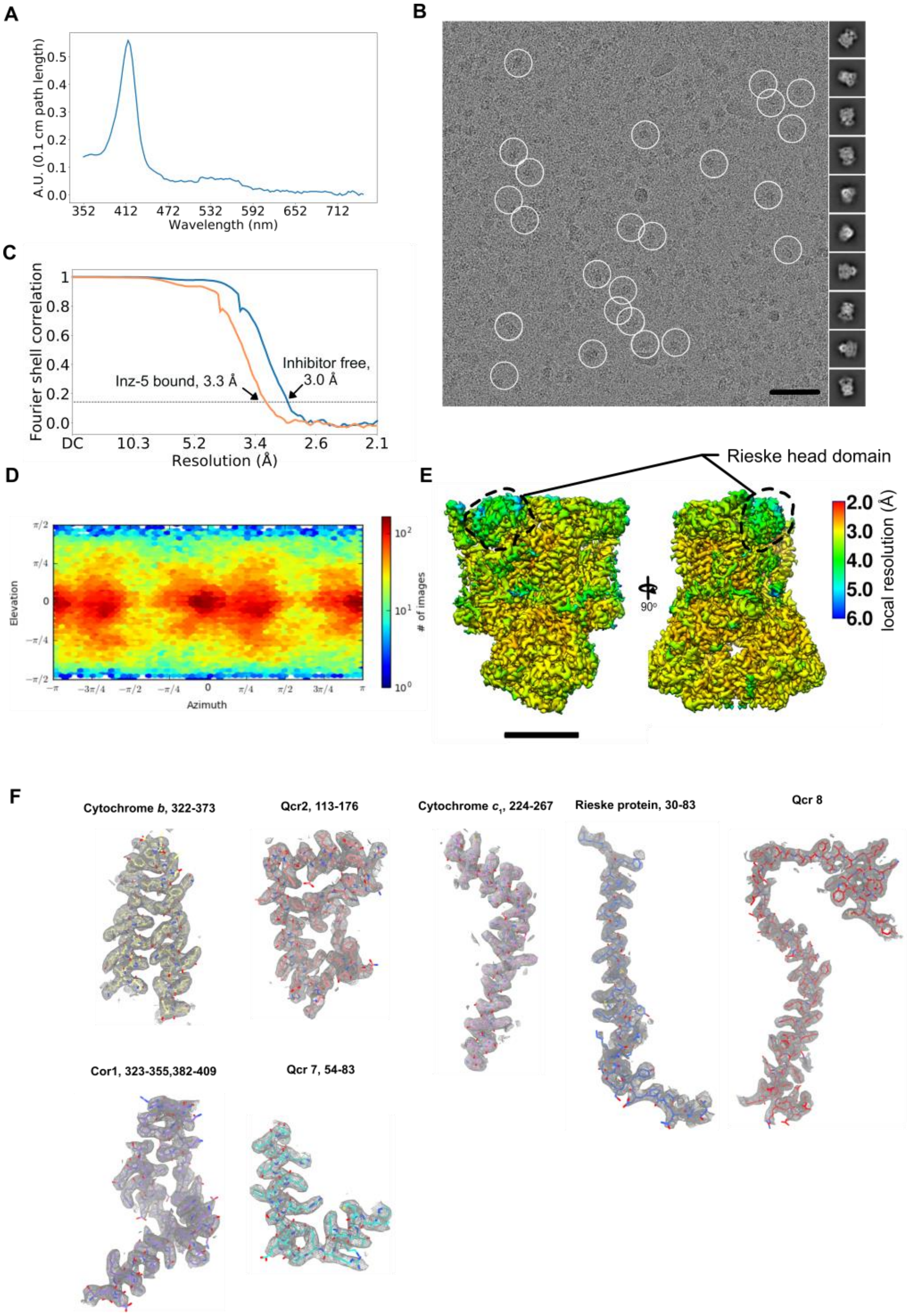


**Fig. S1**. **CryoEM map calculations for inhibitor free dataset.** (**A**) UV-visible spectra of purified $CIII_2$. (**B**) Representative micrograph and 2D class average images for $CIII_2$. Scale bar, 500 Å. (**C**) Fourier shell correlation (FSC) curve after correction for solvent masking. (**D**) Viewing direction distribution for particle images. (**E**) Local resolution estimate for $CIII_2$ map from non-uniform refinement with $C_2$ symmetry. Scale bar, 50 Å. (**F**) Examples of model in map fit.

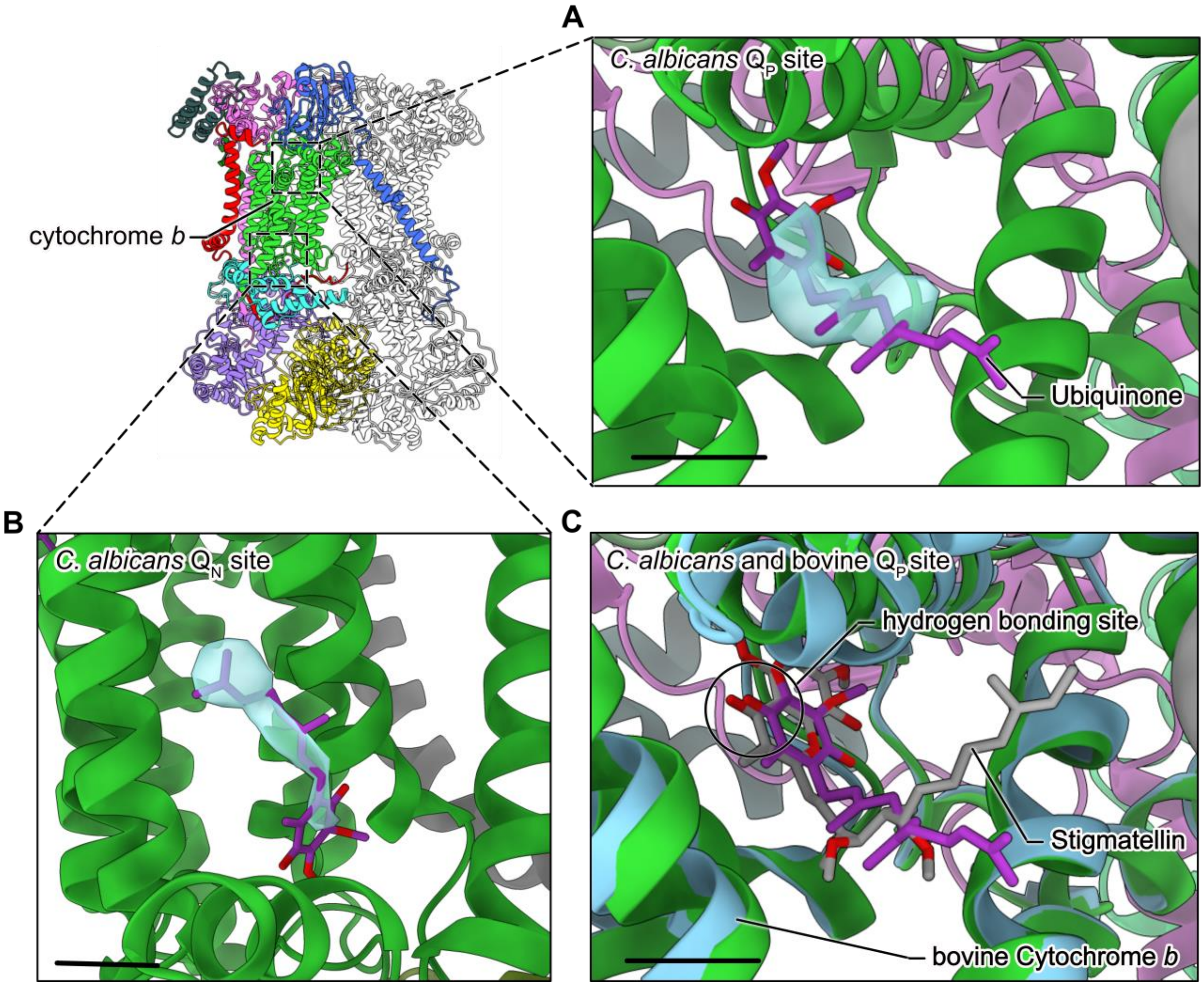


**Fig S2. Endogenous ubiquinone bound to Q$_P$ and Q$_N$ sites.** Ubiquinone from previous structure of bovine $CIII_2$ (Letts *et al.*, 2019) rigid body fit into (**A**) Q$_P$, and (**B**) Q$_N$ sites. (**C**) Overlay of stigmatellin bound bovine cytochrome *b* (Zhang *et al.*, 1998) with current structure. Scale bars, 5 Å.

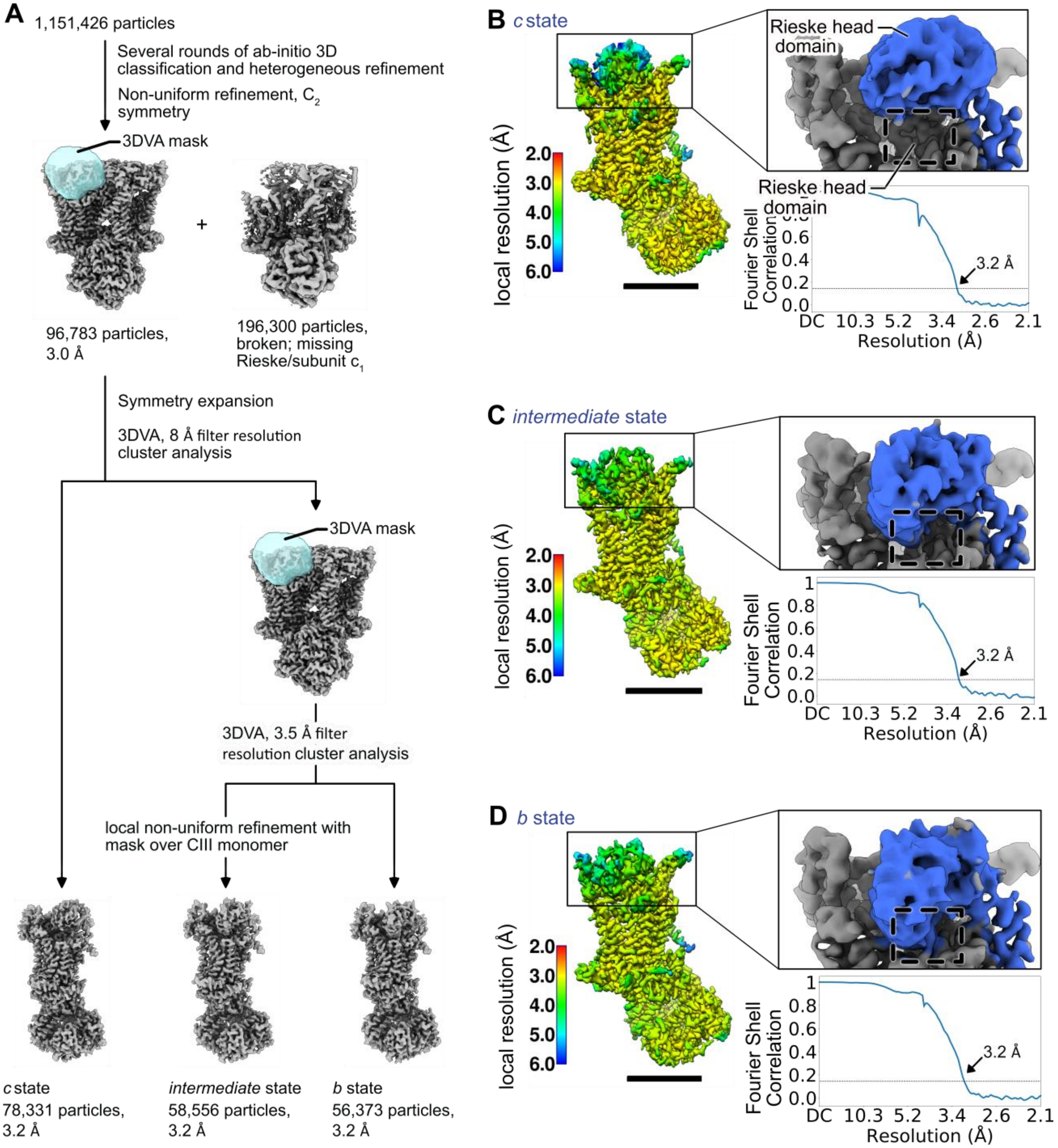


**Fig S3. CryoEM workflow for inhibitor-free dataset.** (**A**) Dataset cleaning and 3DVA workflow. Local resolution estimate, close-up view of Rieske head domain region, and Fourier shell correlation (FSC) curve after correction for solvent masking for local non-uniform refinement of (**B**) *c* state, (**C**) *intermediate* state, and (**D**) *b* state particles. Dashed box in inset highlights conformational changes between the *c*, *intermediate*, and *b* states. Scale bars, 50 Å.

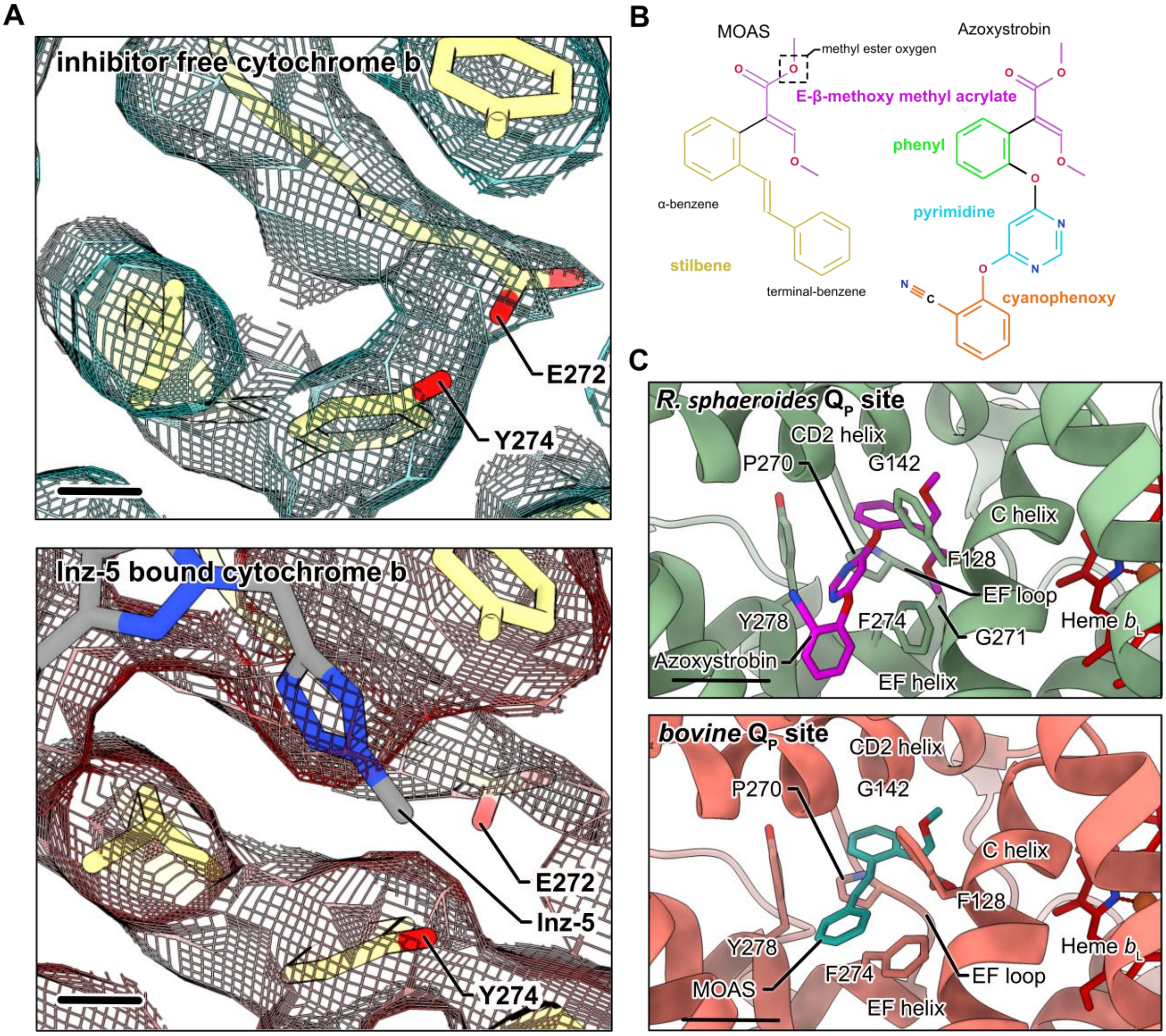


**Fig S4. $Q_P$ binding site conformational changes and strobilurin inhibitor binding poses.** (**A**) CryoEM map and model for $Q_P$ binding site for inhibitor-free (top), and Inz-5-bound dataset (bottom). Scale bars, 2 Å. (**B**) Structure of MOAS and azoxystrobin. (**C**) Atomic model of $Q_P$ site of *R. sphaeroides* CIII with azoxystrobin bound (top)(Esser *et al.*, 2019), and bovine CIII with MOAS bound (bottom)(Esser *et al.*, 2004). Scale bars, 5 Å.

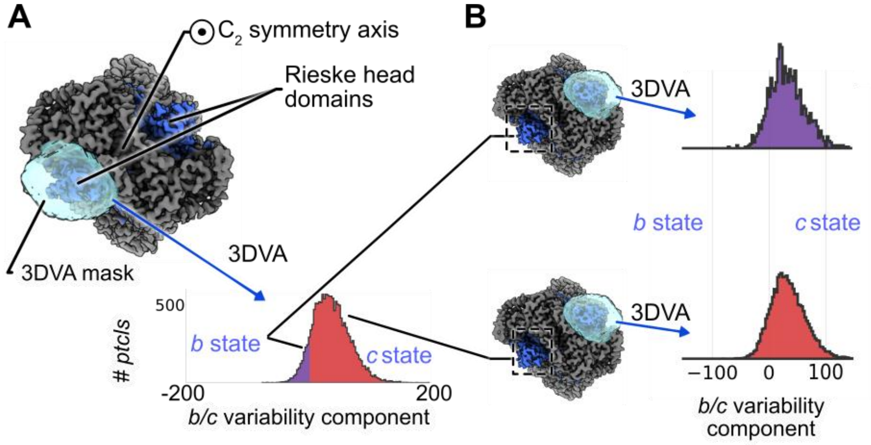


**Fig S5. Assessment of cooperativity between $CIII_2$ Rieske head domains in opposing monomers for Inz-5 bound dataset. (A)** 3DVA of the Rieske head domain from the first monomer was used to divide data into b and c state (purple and red, respectively). **(B)** 3DVA of the Rieske head domain of the second monomer from images where first monomer was in the b state (top) and c state (bottom). There are insufficient particle images for reliable calculation of 3D maps and consequently only variability components are shown.

**References**

Esser, L. *et al.* (2004) 'Crystallographic studies of quinol oxidation site inhibitors: A modified classification of inhibitors for the cytochrome bc1 complex', *Journal of Molecular Biology*, 341(1), pp. 281–302. doi: 10.1016/j.jmb.2004.05.065.

Esser, L. *et al.* (2019) 'Crystal structure of bacterial cytochrome bc1 in complex with azoxystrobin reveals a conformational switch of the Rieske iron–sulfur protein subunit', *Journal of Biological Chemistry*, 294(32), pp. 12007–12019. doi: 10.1074/jbc.RA119.008381.

Letts, J. A. *et al.* (2019) 'Structures of Respiratory Supercomplex I+III2 Reveal Functional and Conformational Crosstalk', *Molecular Cell*. Elsevier Inc., 75(6), pp. 1131-1146.e6. doi: 10.1016/j.molcel.2019.07.022.

Zhang, Z. *et al.* (1998) 'Electron transfer by domain movement in cytochrome bc1', *Nature*, 392(6677), pp. 677–684. doi: 10.1038/33612.
